# Deficiency of the histone lysine demethylase KDM5B causes autism-like phenotypes via increased NMDAR signalling

**DOI:** 10.1101/2024.05.28.596232

**Authors:** Leticia Pérez-Sisqués, Shail U. Bhatt, Angela Caruso, Mohi U. Ahmed, Talia E. Gileadi, Shoshana Spring, Eleanor Hendy, Joyce Taylor-Papadimitriou, Diana Cash, Nicholas Clifton, Jacob Ellegood, Laura C. Andreae, Jason P. Lerch, Maria Luisa Scattoni, K. Peter Giese, Cathy Fernandes, M. Albert Basson

## Abstract

Loss-of-function mutations in genes encoding lysine methyltransferases (KMTs) and demethylases (KDMs) responsible for regulating the trimethylation of histone 3 on lysine 4 (H3K4me3) are associated with neurodevelopmental conditions, including autism spectrum disorder and intellectual disability. To study the specific role of H3K4me3 demethylation, we investigated neurodevelopmental phenotypes in mice without KDM5B demethylase activity. These mice exhibited autism-like behaviours and increased brain size. H3K4me3 levels and the expression of neurodevelopmental genes were increased in the developing *Kdm5b* mutant neocortex. These included elevated expression of *Grin2d*. The *Grin2d* gene product NMDAR2D was increased in synaptosomes isolated from the *Kdm5b*-deficient neocortex and treating mice with the NMDAR antagonist memantine rescued deficits in ultrasonic vocalisations and reduced repetitive digging behaviours. These findings suggest that increased H3K4me3 levels and associated *Grin2d* gene upregulation disrupt brain development and function, leading to socio-communication deficits and repetitive behaviours, and identify a potential therapeutic target for neurodevelopmental disorders associated with KDM5B deficiency.

**Teaser:** Inhibitors targeting NMDA receptors may represent viable therapies for KDM5B neurodevelopmental disorders

## Introduction

Mutations and variants in >50 genes encoding proteins associated with chromatin are linked to neurodevelopmental disorders (*1*). Several of these factors function as regulators of chromatin modified by histone 3 lysine 4 trimethylation (H3K4me3), a post-translational modification typically found at active gene promoters (*2, 3*). As H3K4me3 is permissive for transcription and linked to transcriptional elongation and output (*4*), mutations in genes encoding histone methyltransferases that catalyse H3K4 methylation, or in genes encoding “reader” proteins that mediate downstream effects of H3K4me3, are predicted to be associated with reduced transcription (*5, 6*). Examples include CHD8, an ATP-dependent chromatin remodeller recruited to H3K4me2/3 and KMT2A, a H3K4 methyltransferase. Mutations of the *CHD8* or *KMT2A* genes cause defined syndromes characterised by Autism Spectrum Disorder (ASD) with high penetrance (*7–14*). H3K4me2 levels were found to be reduced in a small sample of brain samples from idiopathic ASD individuals and genetically defined ASD mouse models (*15*). Restoring normal H3K4me2 levels in these mouse models by treatment with H3K4-specific lysine demethylases rescued ASD-associated phenotypes, suggesting that the reduced H3K4me2 levels are responsible for these phenotypes.

Intriguingly, mutations in H3K4me3-specific demethylases of the KDM5 family are also associated with ASD (*12, 13, 16*), suggesting that increased H3K4me3 and transcriptional output could also lead to ASD. A recent forward genetic study identified recessive *Kdm5a* mutations in mice with alterations in ultrasonic vocalisation (USV) emissions and nest building phenotypes and further reported evidence for social, repetitive and cognitive phenotypes in *Kdm5a^-/-^* mice (*17*). Hayek at al. further showed that individuals with recessive and heterozygous *KDM5A* mutations exhibited ASD, developmental delay and intellectual disability. Mutations in the X-linked KDM5C gene that either disrupt its demethylase activity, or keep enzymatic activity intact, cause the intellectual disability Claes-Jenssen syndrome in males (*18, 19*). Recessive mutations in *KDM5B* causes a human syndrome associated with developmental delay, ASD, intellectual disability, hypotonia, craniofacial and limb abnormalities (*20*). In contrast to loss-of-function variants in genes like *CHD8* that are associated with ASD at high penetrance, heterozygous *KDM5B* mutations associated with ASD and developmental delay are not fully penetrant, and are often inherited from apparently unaffected parents (*13, 21–24*). Many ASD-associated missense mutations in *KDM5B* map to the JmjN, ARID and JmjC domains that form the catalytic domain of the protein, suggesting that changes in demethylase activity are associated with the neurodevelopmental phenotypes in these individuals (*21*).

KDM5B is essential for normal development with homozygous loss-of-function mutations in mice resulting in embryonic or significant early postnatal lethality (*24–26*). Similar to Drosophila *Kdm5*, the lysine demethylase activity of KDM5B is not essential for embryonic development as a homozygous mouse model that lacks this activity is viable and fertile with only a subtle mammary gland phenotype reported to date (*26, 27*). These findings are consistent with the observation that H3K4me3 levels only become dysregulated in *Kdm5b*-deficient brain after E17.5 (*25*). The reason why *Kdm5b* deficiency has little effect on H3K4me3 levels in the brain in the mid-gestation embryo is not known, but the most likely possibility is compensation by other KDM5 family members.

In addition to mutations in genes encoding chromatin remodelling factors and synaptic proteins, de novo variants in genes encoding N-methyl-D-aspartate receptor (NMDAR) subunits have been identified in ASD probands and both reduced and elevated NMDAR activity have been implicated in ASD (*28, 29*). NMDARs are ligand-gated cation channels that mediate excitatory synaptic transmission (*30*). Increased NMDAR function during the early postnatal period (prior to P21), has been reported in specific ASD rodent models, including a *Shank2^e6-7^* mutant and valproic acid exposure model (*31, 32*). Treatment of *Shank2^e6-7^*mutant mice at the time of NMDAR overactivity (P7-P21) with a non-competitive NMDAR antagonist, memantine (*33*), rescued the sociability phenotype (*32*). Memantine treatment of adult *IRSp53* mutant mice was also able to restore social interactions but not a hyperactivity or adult ultrasonic vocalisation phenotype (*34*). The NMDAR antagonist agmatine rescued the social, hyperactivity and repetitive behavioural phenotypes in a VPA-exposed rat ASD model (*35*). Intriguingly, drugs that target the epigenome have been shown to restore NMDAR function in *Shank3* mutant mice (*36–38*), further linking the epigenome with NMDAR dysfunction in specific ASD subtypes.

To test if dysregulation of the H3K4me3 demethylase activity of KDM5B is sufficient to cause ASD, we characterised mice that lack KDM5B demethylase activity (*27*). These mice were found to exhibit ASD-like behavioural phenotypes. Molecular studies revealed increased H3K4me3 levels and gene expression abnormalities in the early postnatal brain. These changes included an upregulation of the NMDAR2D subunit, encoded by the *Grin2d* gene. Treatment with memantine increased ultrasonic vocalisation emission in pups and reduced excessive digging behaviour in adult mice, suggesting that NMDAR hyperactivation may be responsible for ASD-like behavioural phenotypes.

## Results

### KDM5B is a H3K4me3 demethylase in the developing neocortex

To study the function of the histone demethylase KDM5B and avoid complications due to embryonic or early postnatal lethality, we established a mouse line in which exons 2 to 4 were deleted, resulting in the truncation of the carboxyl end of the JmjN domain and deletion of the entire ARID domain (Fig. 1A). This mutation specifically disrupts the H3K4me3 demethylase activity of the protein, leaving the developmental functions of KDM5B required for embryonic development and postnatal survival intact (*26, 27*). Albert et al. have shown that *Kdm5b* deletion only affects H3K4me3 levels from late embryonic brain development (*25*), so we focussed our analysis on postnatal stages.

**Fig. 1.**
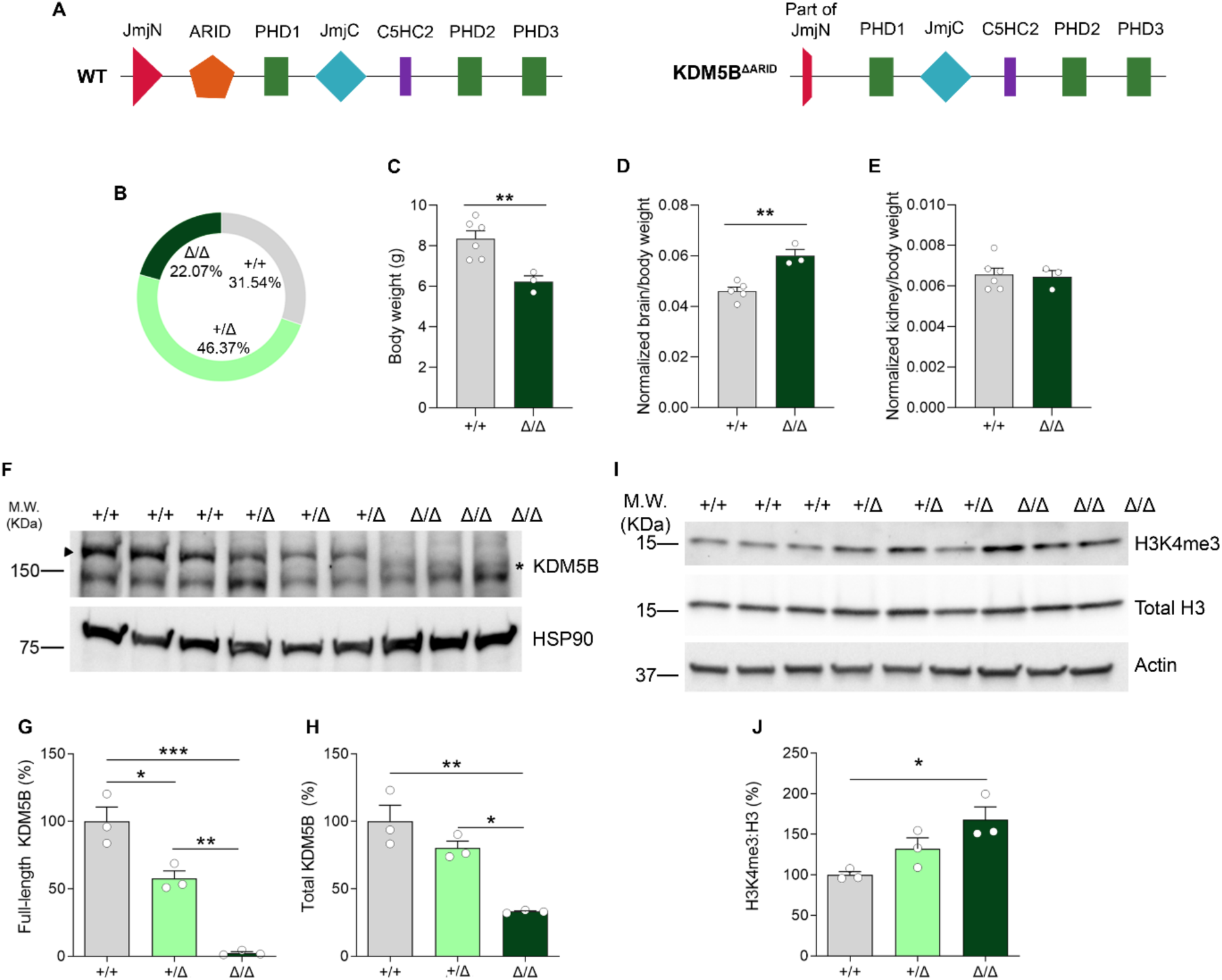
Developmental alterations and KDM5B expression in control and ΔARID mice. **A)** Schematic representations comparing mouse WT and ΔARID KDM5B linear domain structures. Note that transgenic *Kdm5b ΔARID* mice carry a deletion of the entire ARID domain and a truncation of the carboxyl end of the JmJ domain. **B)** Genotyping frequencies show no statistically significant differences between genotypes (Chi-square test) at postnatal day (PND) 21. **C-E)** *Kdm5b ΔARID* homozygous mice (*Kdm5b*^Δ/Δ^) exhibit decreased weight and increased brain-to-body weight ratios, suggesting developmental defects. Brain and kidney weights did not differ between genotypes (data not shown; One way-ANOVA for brain weight, p=0.9323; One-way ANOVA for liver weight, p=0.8144). **F-H)** Estimation of full-length (G) and total (H; full length and truncated) KDM5B protein expression by western blot in neocortical samples extracted from P5 animals. HSP90 was used as loading control. Arrowhead: Full-length KDM5B isoform; asterisk: truncated ΔARID protein. **I,J)** H3K4me3 and total H3 levels in WT, heterozygous and ΔARID homozygous mice in neocortical samples extracted from P5 animals. Wild type (+/+) and ΔARID heterozygous (+/Δ) and homozygous (Δ/Δ) mice. Data is shown as percentage versus WT average. Data is shown as mean ±SEM and was analysed with One-way ANOVA followed by Tukey’s multiple comparisons test. *p<0.05, **p<0.01 and ***p<0.001.

Homozygous *Kdm5b^ΔARID/ΔARID^* (*Kdm5b^Δ/Δ^*) mice on a C57BL/6J background were born within expected ratios from *KDM5B^+/Δ^* intercrosses (+/+: 31.54%; +/Δ: 46.37%; Δ/Δ: 22.07%, n=69, χ^2^ test, p=0.6077) with no reduction in postnatal survival (Fig. 1B). When examined at postnatal day 12 (PND 12), homozygous mutant mice displayed decreased body weight compared to wildtype littermates, suggesting some effect on their ability to grow or thrive (Fig. 1C). Intriguingly, at this stage mutant mice exhibited increased brain-to-body weight ratios (Fig. 1D), whilst the relative size of other vital organs was not affected (Fig. 1E), suggesting specific neurodevelopmental functions for KDM5B.

To determine the effects of the ΔARID mutation on KDM5B protein in the brain, we visualised KDM5B by immunoblot in brain lysates. The shorter KDM5B ΔARID protein was detected in hippocampal and neocortical samples from postnatal day 5 (PND5) *Kdm5b^+/Δ^* and *Kdm5b^Δ/Δ^* pups, with the expected reduced molecular weight. In line with previous results (*27*), full length KDM5B protein was reduced by 50% in the neocortex of *Kdm5b^+/Δ^*mice and absent in *Kdm5b^Δ/Δ^* mice (Fig. 1F,G). Importantly, total (full-length plus ΔARID) KDM5B protein levels were reduced in the neocortex of homozygous mice compared to wildtype littermates (Fig. 1F,H). Thus, in addition to the reported lack of demethylase activity, our data suggest that the ΔARID protein is also slightly less stable than the full-length protein, although the possibility that the antibody is less efficient at detecting the mutant protein cannot be excluded.

The ΔARID protein lacks demethylase activity in in vitro assays (*27*). To determine if KDM5B functions as a H3K4me3 demethylase in the developing brain, and if steady-state H3K4me3 levels are altered in *KDM5B^Δ/Δ^* mice, we compared H3K4me3 levels in brain tissue from wildtype, heterozygous and homozygous mice. We detected a significant increase in H3K4me3 levels in neocortical samples from homozygous, but not heterozygous mice (Fig. 1I,J).

### Autism-associated behavioural phenotypes in *Kdm5b^Δ/Δ^* mice

To determine if a reduction in KDM5B levels and lack of KDM5B demethylase activity are sufficient to cause behavioural phenotypes in mice, we assessed socio-communicative, repetitive, anxiety and motor behaviours (Fig 2A). Both male and female animals were tested (Sup. Fig. 1), and data were combined when no significant sex differences were observed (Fig. 2). Homozygous pups emitted significantly fewer USVs upon separation from the mother (Fig. 2B). Homozygous adults exhibited no significant differences in social approach and social investigation compared to wildtype littermates in the social investigation and 3-chamber and tests, respectively (Fig. 2C,D). In the marble burying test, a measure of repetitive behaviours, mutant mice buried marbles significantly faster than wildtype littermates, indicative of compulsive, repetitive digging behaviours (Fig. 2E). General locomotion and anxiety levels did not differ between genotypes in an open field arena (Fig. 2F) (Suppl. Fig. 1E,F). As further confirmation, no differences were found in the elevated plus maze test (EPM) (Fig. 2G), another test for anxiety. Homozygous mice showed reduced performance compared to wildtype animals in the accelerating rotarod task (Fig. 2H). *Kdm5b^Δ/Δ^* mice exhibited reduced hindlimb grip strength (Fig. 2I,J), which likely contributed to their poor performance on the rotarod test and is consistent with a report of hypotonia in an individual with recessive *KDM5B* mutations (*39*).

**Fig. 2.**
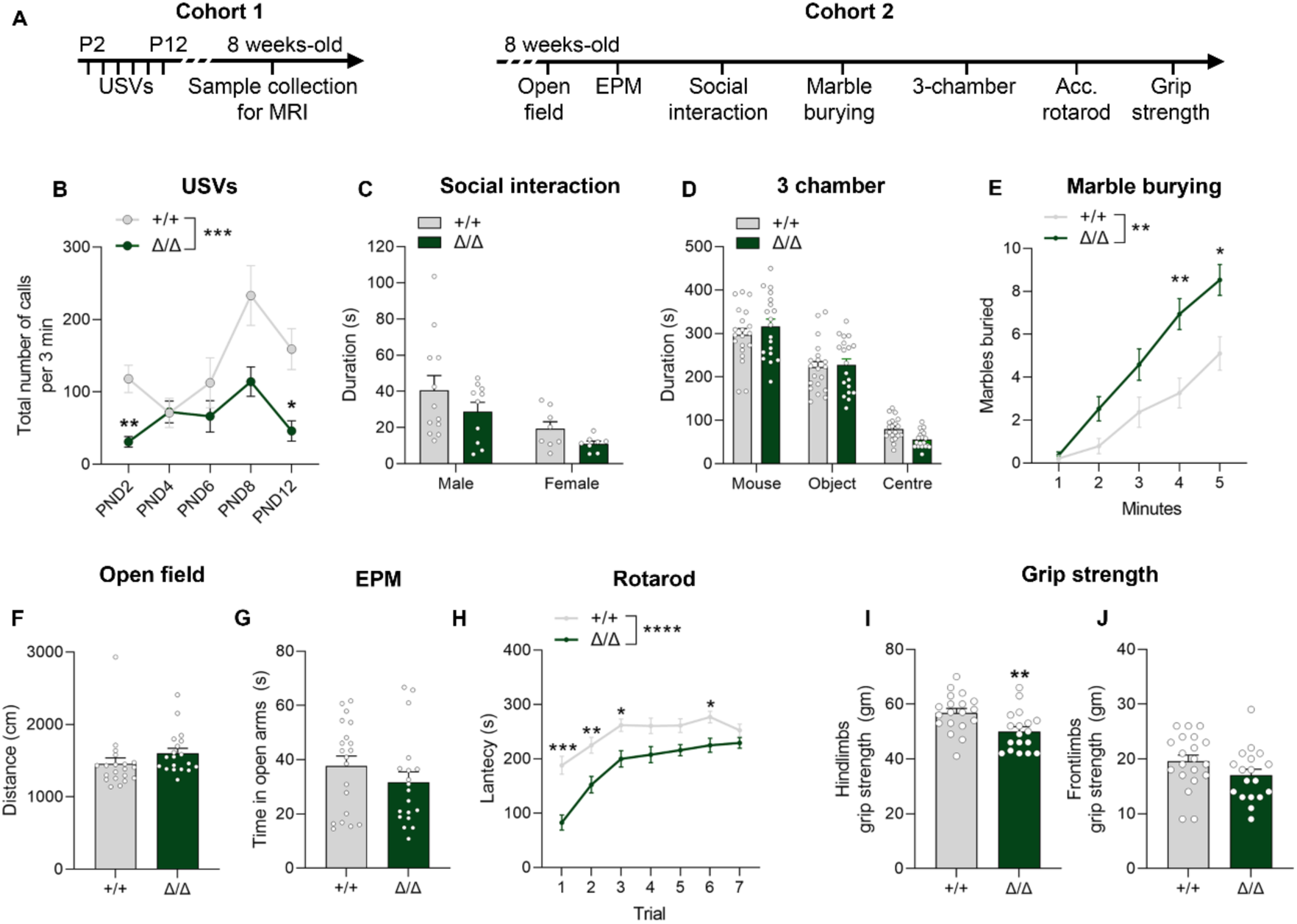
Behavioural assessment of *Kdm5b*^Δ/Δ^ mice. Behavioural assessment of a cohort of neonatal (B) (+/+ n=3 males and 7 females; Δ/Δ n=6 males and 6 females) and adult mice (C-J) (+/+ n=11-12 males and 7-8 females; Δ/Δ n=10-11 males and 6-8 females). **A)** Experimental time-course. Note that different cohorts were used for experiments with neonatal and adult mice. **B)** Total number of ultrasonic vocalisations (USVs) during 3 minutes testing sessions on indicated postnatal days. Repeated measures two-way ANOVA Interaction: F(_4,80_)=2.866, *p=0.0284; Genotype effect: F_(1,20)_=15.03, ***p=0.0009. **C)** Duration, in seconds, of social investigation, defined as total duration of sniffing above the shoulders of a conspecific mouse. Two-way ANOVA sex effect: F_(1,34)_=9.785, **p=0.0036. **D)** Time spent, in seconds, in each chamber in the 3 chamber sociability test. **E)** Number of marbles buried during a 5 minute test. Repeated measures two-way ANOVA Interaction: F_(4, 136)_=5.894, ****p=0.0002; Genotype effect: F_(1,34)_=11.08, **p=0.0021. **F)** Distance travelled in the outer area of an open field arena during a 5 minute test. **G)** Time spent in the open arms of the elevated plus maze (EPM) is shown. **H)** Mean latency of mice to fall from the rotarod during 7 trials in one day. Repeated measures two-way ANOVA Interaction: F_(6,222)_=2.944, **p=0.0088; Genotype effect: F_(1,37)_=21.54, ****p<0.0001. **I,J)** Hind- and frontlimb grip strength measurements. Data is shown as mean ±SEM and was analysed with Two-way ANOVA (C,D), repeated measures ANOVA (B,E,H) followed by Sidak’s post hoc test, and Student’s t-test (F,G,I,J). \**P*<0.05, \*\**P*<0.01 and \*\*\**P*<0.001.

### Structural brain anomalies associated with KDM5B deficiency

Intact brains were collected from the mice that underwent ultrasonic vocalisation phenotyping (Fig. 2A). Structural magnetic resonance imaging (MRI) was performed to identify changes in absolute and relative volumes. We assessed 290 brain regions with divisions across the cortex, subcortical areas, cerebellum, brain stem, ventricular systems and fibre tracts (see Supplementary tables 1-4 for full results tables). A 4.2% (±1.06%) increase in brain volume was detected in homozygous mutant animals (Fig. 3A). Moreover, absolute volumes for the fibre tracts and the ventricular systems were also increased in *Kdm5b^Δ/Δ^* mice (6.02±0.62% and 15.23±10.47%, respectively) (Fig. 3B,C). Alterations within the fibre tracts included the anterior commissure (temporal limb) and the corticospinal tract. Mutant mice showed increases in volume across cortical areas such as the anterior olfactory nucleus, the endopiriform nucleus, the taenia tecta, the infralimbic area, the primary somatosensory area associated to the nose, and the dorsal auditory area (11.28±1.9%, 8.57±2.04%, 8.84±1.56%, 11.17±0.36%, 7.39±3.06% and 9.34±0.61% increases, respectively) (Supplementary tables 1 and 3). *Kdm5b^Δ/Δ^*mice also showed robust increases in volume in cerebral nuclei such as the striatum (+7.52±1.52%), dorsal pallidum (+9.04±3.38%), lateral septum (+7.32±0.43%), the thalamus (+5.54±1.77%), and in the cerebellar nodular lobe (+15.67±2.07%).

**Fig. 3.**
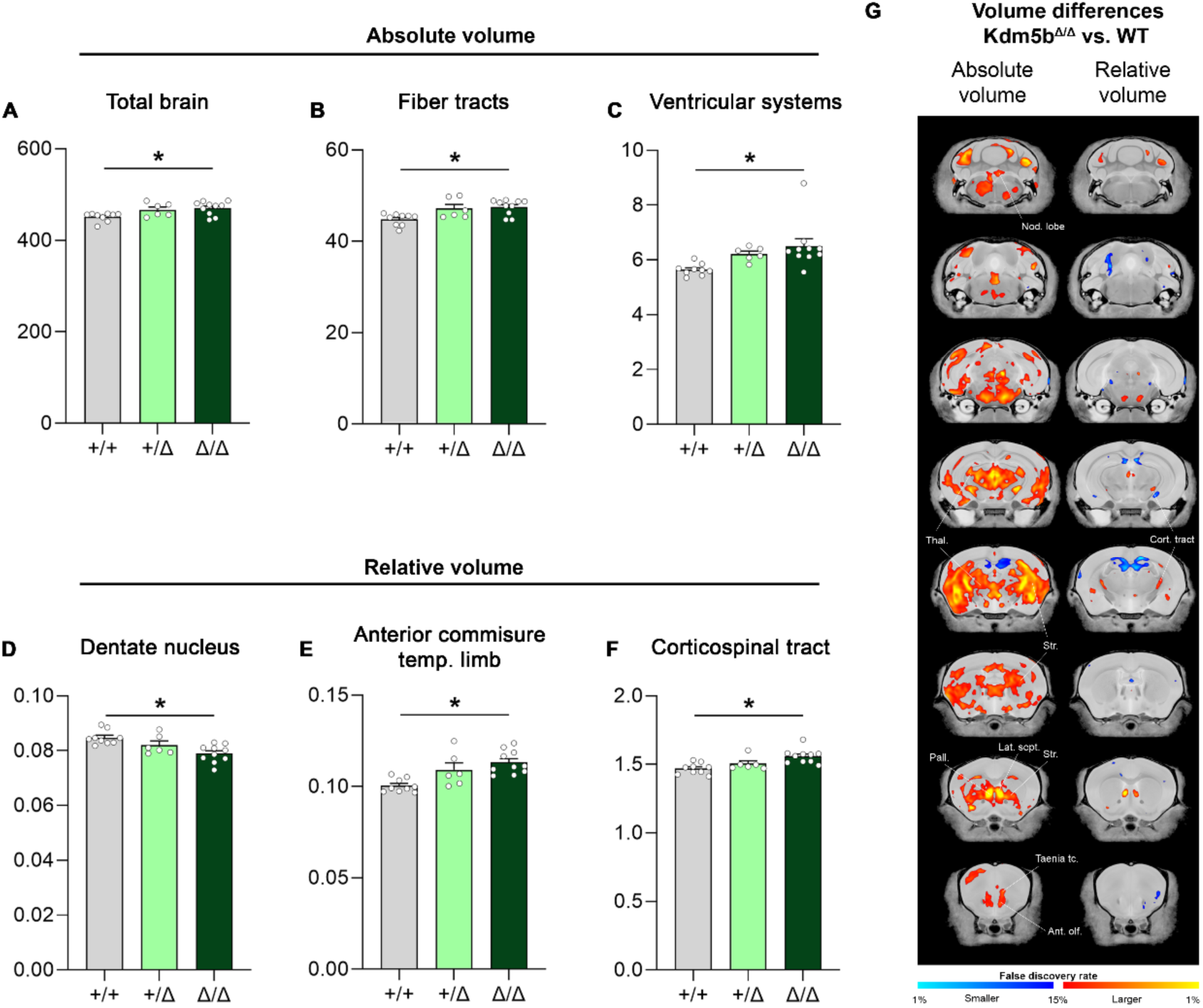
*Kdm5b*^Δ/Δ^ mutant mice display megalencephaly and altered regional volume of different brain areas. **A-C)** MRI analysis revealed significant increases in absolute brain size together with increased regional volume of different brain areas such as the fibre tracts and the ventricular systems. **D-F)** Correction for absolute brain volume revealed significant changes in volume in the dentate nucleus, corticospinal tract and anterior commissure of the temporal limb. **G)** Voxel-wise differences in volume (absolute, left; relative, right) between wild-type and *Kdm5b*^Δ/Δ^ mutant littermates. Data is shown as mean ±SEM. Multiple comparisons (all brain regions) were controlled for using the False Discovery Rate (see Supplementary tables 1-4). *FDR<0.05.

After correction for total brain volume, relative volumes were significantly reduced in the dentate nucleus (−6.84±0.9%) but still significantly larger in the anterior commissure of the temporal limb (+12.64±2.18%) and in the corticospinal tract (+6.16±0.68%) (Fig. 3D-F). Voxel-wise differences showed similar trends (Fig. 3G). We did not detect significant differences between wild-type and heterozygous animals in either absolute or relative volumes, although similar trends were apparent compared to homozygotes (Sup. Fig. 2 and Supplementary tables 2 and 4). Accordingly, brain weight negatively correlated with wildtype *Kdm5b* copy number (Sup. Fig. 2B-G).

### KDM5B regulates the expression of neurodevelopmental genes

The increased H3K4me3 observed in the PND5 neocortex of *Kdm5b*^Δ/Δ^ mice (Fig. 1G) is predicted to result in increased gene expression. To test this hypothesis, gene expression in *Kdm5b*^Δ/Δ^ and wildtype PND5 neocortical and hippocampal tissues were compared by RNA sequencing (RNA-seq) (Supplementary tables 5 and 6). This analysis identified 177 differentially expressed genes (DEGs, FDR<0.05) in the neocortex (Fig. 4A) and 162 DEGs in the hippocampus (Fig. 4F). The transcriptomic differences between the two genotypes were visualised in Volcano plots and heatmaps, where the majority of transcripts were upregulated in *Kdm5b^Δ/Δ^* mice, both in the neocortex (Fig. 4A,B) (88%) and hippocampus (Fig. 4F,G) (85%), in support of our hypothesis.

**Fig. 4.**
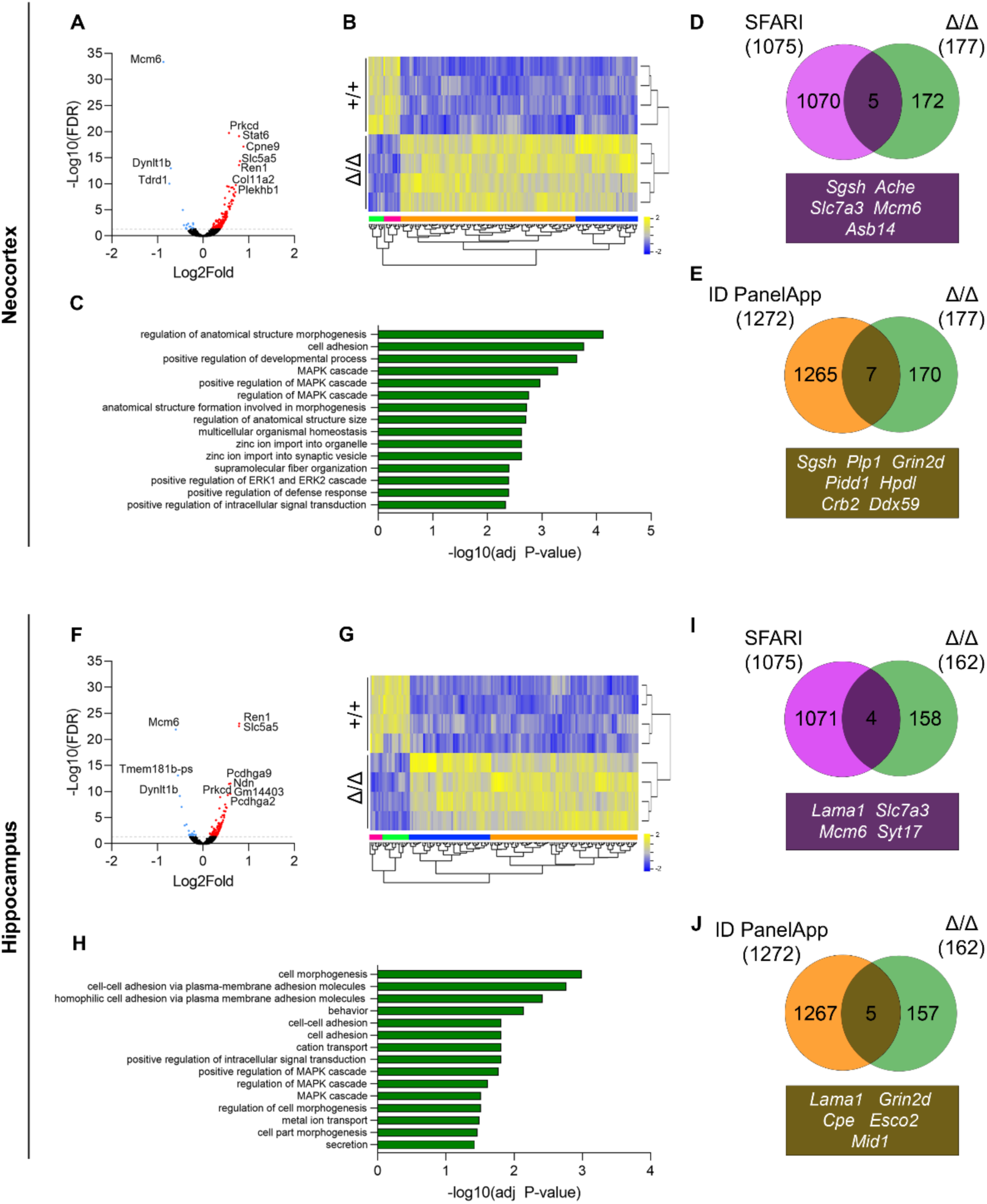
Gene expression changes in PND5 *Kdm5b*^Δ/Δ^ neocortex and hippocampus determined by RNAseq (n=4 mice per genotype, 2 males and 2 females). **A, F)** Volcano plot displaying gene expression changes detected by DESeq2 in neocortex (A) and hippocampus (G). Each point represents an individual gene, and all DEGs (FDR<0.05) are highlighted in red (upregulated) or blue (downregulated). The top 10 differentially expressed genes are depicted next to their corresponding position. **B, G)** Heatmap of sequenced genes in neocortical (B) and hippocampal (H) samples. Yellow, upregulated; blue, downregulated. **C, H)** Gene ontology for the DEGs (FDR<0.05) in the neocortex (C) and hippocampus (I). The ten most significant “GO Biological pathways” hits are shown for each set (term size < 1500). **D, I)** Venn diagrams showing the extent of overlap between DEGs (FDR<0.05), and ASD-associated genes obtained from the SFARI human Gene database (accessed September 2022). The overlapping genes are shown below the diagram. **E, J)** Venn diagrams showing the extent of overlap between DEGs (FDR<0.05), and ID-associated genes obtained from the Genomics England PanelApp Intellectual disability database (version 4.42, accessed December 2022). The overlapping genes are shown below the diagram.

Gene ontology analyses were performed to identify potential biological pathways affected in *Kdm5b*^Δ/Δ^ mutants (Supplementary tables 7 and 8). This analysis identified a significant enrichment of developmental, signalling and metal ion transport-associated genes (Fig. 4C,H). A few differentially expressed genes were known SFARI ASD-associated genes (Fig. 3D,I), and ID PanelApp intellectual disability-associated genes (Fig. 3E,J) (Supplementary tables 9 and 10). These ASD/ID-associated genes included the most significantly downregulated gene, *Mcm6* (*12, 13, 40*), the downregulated gene *Slc7a3*, which encodes the Cationic amino acid transporter CAT3 (*41*), and the NMDAR subunit gene *Grin2d*, which was upregulated.

### Treatment with an NMDAR antagonist can improve behavioural deficits

As KDM5B is expected to function as a repressor, genes upregulated in *Kdm5b* mutants are most likely directly regulated by KDM5B. One of the most interesting neurodevelopmental genes upregulated in these mutants was *Grin2d*, which encodes an NMDAR subunit (NMDAR2D). Microduplications at 19q13.33 that includes the *GRIN2D* gene (*42*), and both de novo gain- and loss-of-function mutations in *GRIN2D* are associated with neurodevelopmental disorders (*43*),(*44*). Bi-directional changes in NMDAR signalling have also been implicated in ASD (*28*).

First, to determine if the NMDAR2D subunit, encoded by the *Grin2d* gene, was indeed increased in the PND5 neocortex of *Kdm5b*^Δ/Δ^ mutants, we quantified NMDAR2D protein levels in purified synaptosomes by Western blot. This analysis confirmed increased NMDAR2D levels in mutant synaptosomes, compared to wildtype controls (Fig. 5A,B). To determine if inhibition of NMDARs could rescue some of the neurodevelopmental phenotypes associated with KDM5B deficiency, we treated mice with the NMDAR antagonist memantine and examined the effect on USV and repetitive digging behaviours. Memantine or saline were administered intraperitoneally 30 minutes before the start of each test (Fig. 5C). The doses were 5.6mg/kg for USVs and 10mg/kg for repetitive and motor learning behaviours. Memantine significantly increased the number of ultrasonic calls emitted following maternal separation in mutant mice (repeated measures Two-way ANOVA interaction effect F(_4,48_)=3.520, *p=0.0134), most notably at postnatal day 8 (Fig. 5D), when the number of calls peaked in the previous experiment (see Fig. 2B). However, the number of USVs was not significantly affected in wild-type animals (Fig. 5E), suggesting memantine is targeting a disease-specific mechanism in mutant mice. Memantine treatment of adult mice strongly suppressed the repetitive digging behaviour in both the *Kdm5b*^Δ/Δ^ mutant mice and wild-type littermates (Fig. 5F,G), therefore likely targeting a mechanism not specific to *Kdm5b* deficiency. An acute administration of the drug did not improve the performance in the accelerating rotarod task. Intriguingly, while memantine reduced the latency in wild-type animals (Fig. 5I), it did not significantly affect the performance of mutant mice (Fig. 5H). These findings suggest NMDAR antagonist treatment as a potential therapy for socio-communicative deficits and repetitive symptoms associated with *KDM5B* deficiency.

**Fig. 5.**
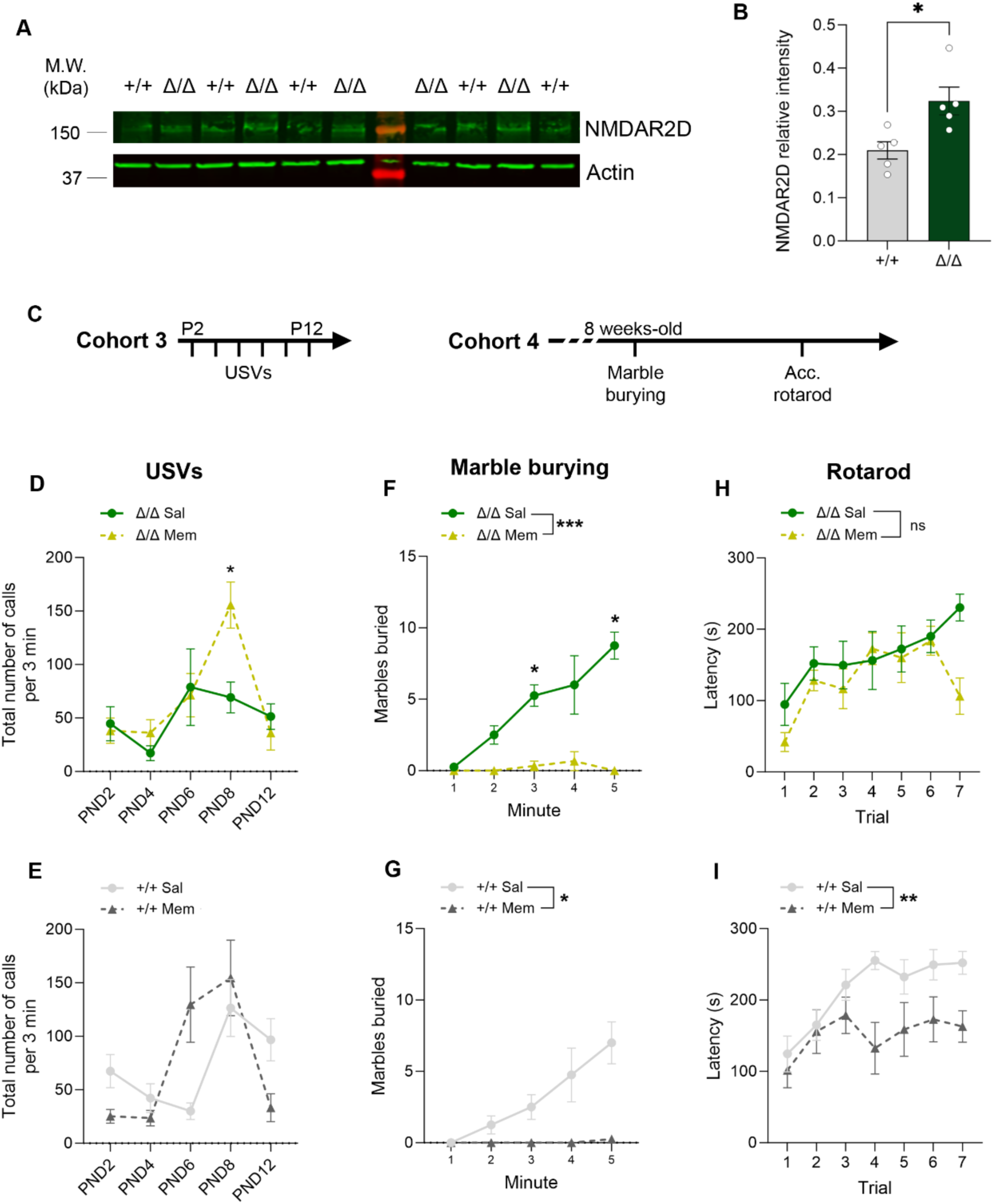
Memantine ameliorates neonatal communication deficits and digging behaviour in *Kdm5b* mutant mice. **A)** Immunoblotting for NMDAR2D (green) in synaptosomes extracted from neocortical samples obtained from P5 wild-type and mutant animals. β-actin (green) was used as a loading control protein ladder (red) and molecular weights are also shown. **B)** Densitometric quantification of NMDAR2D protein expression as in A). **C)** Experiment design. Two different cohorts were used for neonatal and adult tests. Memantine was administered 30 minutes before the start of the test each day. **D,E)** Total number of ultrasonic vocalisations (USVs) during 3 minutes testing sessions on indicated postnatal days. Repeated measures two-way ANOVA Interaction in D: F_(4,48)_=3.520, *p=0.0313 followed by Sidak’s multiple comparisons test (PND8, *p=0.0313). N= 15 +/+ saline, 13 +/+ memantine, 6 Δ/Δ saline and 8 Δ/Δ memantine. **F,G)** Number of marbles buried during a 5-minute test with 2-months-old mice. Repeated measures Two-way ANOVA treatment effect: F_(1,6)_=13.56, *p=0.0103 (G) and F_(1,5)_=99.5, ***p=0.0002 (F) and interaction effect F_(4,24)_=8.387, ***p=0.0002 (G) and F_(4,204)_=5.12, **p=0.0053 (F). N= 4 +/+ saline, 4 +/+ memantine, 4 Δ/Δ saline and 3 Δ/Δ memantine. **H,I)** Mean latency of mice to fall from the rotarod during 7 trials in one day. Repeated measures Two-way ANOVA treatment effect: F_(1,14)_=10.48, **p=0.006 (I) and F_(1,11)_=1.996, p=0.1854 (H) and interaction effect F_(6,84)_=1.524, p=0.1803 (I) and F_(6,66)_=1.9, p=0.0938 (H). N= 10 +/+ saline, 6 +/+ memantine, 7 Δ/Δ saline and 6 Δ/Δ memantine. Memantine doses were 5.6mg/kg for neonatal animals in the USVs test and 10mg/kg for adult mice. Memantine or saline were administered 30min before the start of the test. Data is shown as mean ±SEM.

## Discussion

In this study, we characterised the neurodevelopmental phenotypes of a mouse model homozygous for a hypomorphic, demethylase-deficient allele of *Kdm5b* on a C57BL/6J genetic background. These mice exhibited ASD-associated socio-communicative, repetitive and motor phenotypes. Steady-state H3K4me3 levels were elevated and a number of genes were differentially expressed in the early postnatal neocortex of these mice, compared to wildtype littermates. As expected for a transcriptional repressor and consistent with the increased H3K4me3 levels, the majority of differentially expressed genes were upregulated. As one of these upregulated genes, *Grin2d*, encodes an NMDAR subunit implicated in neurodevelopmental disorders, we treated mice with the NMDAR antagonist, memantine, and found that this treatment ameliorated socio-communicative deficits in pups and repetitive behaviours in adults. Together, our findings show that KDM5B deficiency results in neurodevelopmental phenotypes, suggest a potential mechanistic link with NMDAR gain-of-function conditions and suggest that NMDAR antagonists may represent a potential therapeutic avenue for improving socio-communicative function in individuals with *KDM5B* loss-of-function variants.

The ΔARID mouse model was engineered to delete part of the JmjN and the whole ARID domain to generate a KDM5B protein that lacks demethylase activity (*26*). A recent study confirmed that this so-called ΔARID protein lacks demethylase activity, most likely due to structural protein alterations in the catalytic Jmj domains (*27*). Here, we presented data to suggest that the ΔARID protein is present at reduced levels in the brain compared to the wildtype, full-length protein. As previously reported, homozygous ΔARID mice were viable, and we found no evidence for reduced viability (*26*). This contrasts with other homozygous *Kdm5b* loss-of-function mouse models that exhibited pronounced reductions in embryonic or early postnatal viability (*24–26*). Intriguingly, the homozygous null *Kdm5b* mice with an exon 7 deletion on a C57BL/6NJ background that survive to adulthood, were reported to have a subtle reduction in social interactions and a deficit in social memory (*24, 26*). Together, these findings strongly implicate KDM5B deficiency in behavioural phenotypes relevant to ASD.

The initial report of de novo “likely gene disrupting” and missense variants in *KDM5B* in probands with ASD in simplex families also reported similar variants in the unaffected control population (*12*). It therefore remained unclear if, and to what extent these variants contribute to neurodevelopmental phenotypes. A number of recent studies has now confirmed beyond any doubt that *KDM5B* variants are associated with these neurodevelopmental changes, but that these effects are incompletely penetrant. As a consequence, ASD-associated *KDM5B* variants are amongst the most common inherited ASD-causing variants identified to date (*13, 23*). The identification of genetic modifiers of *KDM5B* penetrance and expressivity will be of significant interest. Our findings suggest that variants affecting the NMDAR pathway may be potent modifiers of the social and repetitive phenotypes.

KDM5B genotype-phenotype correlations in the context of human neurodevelopmental disorders will be of considerable interest given our findings from the demethylase-deficient mouse strain. A recessive syndrome with compound heterozygosity for two different *KDM5B* variants, which included protein-truncating, splice site and missense mutations in the JmjC domain, have been reported (*20, 39*).

To what extent a reduction in H3K4me3 demethylation contribute to this condition remains unknown. Moreover, Wang et al. reported an apparent enrichment of heterozygous missense mutations in and around the demethylase domains (JmjN, ARID and JmjC) of *KDM5B* in individuals with ASD and developmental delay (*21*), implicating demethylase activity in these conditions. However, other studies suggest that most of the likely damaging variants in *KDM5B* are found around the PHD3-PHD3 region. KDM5B interacts with other transcription factors through its PLU domain (*45*), and with HDAC4 through its N-terminal PHD regions (*46*). Further studies are necessary to decipher the potential impact of different *KDM5B* variants on specific mechanisms, including those that may be independent from its demethylase activity. Based on our findings, we propose that variants altering the demethylase activity of KDM5B would primarily affect neurodevelopment postnatally (e.g. via NMDAR signalling), while demethylase-independent variants may affect developmental processes such as neuronal specification and differentiation in the early embryo.

Human patients carrying *KDM5B* variants display a varying degree of brain structural alterations, ranging from a normal MRI to agenesis of corpus callosum, cerebellar vermis alterations and megalencephaly (*20, 24*). This suggests that the disorder could be also associated with brain morphology alterations, although this possibility has yet to be robustly studied, as MRI analyses are not available for all patients. Notably, one patient showed megalencephaly (*24*), in line with the general increase in absolute brain volumes observed in our *Kdm5b^Δ/Δ^*model. Strikingly, we found increased absolute and relative volumes in fibre tracts such as the corticospinal tract and the anterior commissure (temporal). We did not observe alterations specific to the corpus callosum, whereas four individuals with recessive *KDM5B* variants have shown agenesis of the corpus callosum (*39*). This discrepancy suggests that corpus callosum agenesis is not caused by alterations in the demethylase activity of KDM5B, but other, independent KDM5B functions. Interestingly, three individuals with recessive *KDM5B* variants exhibited altered/unsteady gait (*24*). This phenotype could be related to the structural changes we observe in the cerebellum in *Kdm5b^Δ/Δ^* animals. The thalamus and brain stem, two crucial areas controlling posture and gait, are also showing increased absolute volumes in *Kdm5b^Δ/Δ^* homozygous mice. The aberrant size of the striatum is in line with the reduced performance of *Kdm5b^Δ/Δ^* mice in the accelerating rotarod task (*47*). De novo, likely gene disrupting variants in *KDM5B* have also been linked to primary motor stereotypies, another phenotype associated with alterations in the cerebellum (*48*) and cortico-striatal-thalamo-cortical pathways (*49*),(*50–52*). Abnormalities in these brain regions and pathways could underlie the repetitive behaviours observed in *Kdm5b^Δ/Δ^*mice.

Chromatin modifiers and remodellers typically regulate hundreds of gene loci throughout the genome, and function in multiple cell types and at different stages of development. For instance, in the case of CHD8, many ASD-associated genes are dysregulated in the *Chd8*-deficient brain, making it difficult to determine the exact downstream genes responsible for neurodevelopmental phenotypes (*53–58*). We were therefore surprised to find functional evidence that treatment with an NMDAR antagonist was sufficient to improve some for the ASD-associated behavioural phenotypes in our homozygous *Kdm5b* mutants. Memantine is a non-competitive antagonist of NMDARs with low affinity and rapid kinetics (*59*). This treatment presumably normalises an abnormal gain in NMDAR function due to *Grin2d* upregulation in the early postnatal brain. *Grin2d* expression in rodents starts at late embryonic stages, peaking at PND7 (*60*). In the hippocampus NMDAR2D-containing receptors mediate synaptic transmission in interneurons(*61*) and they constrain short-term and long-term potentiation in area CA1 (*62*). In humans, gain-of-function mutations in the NMDAR subunit gene *GRIN2D* are associated with GRIN2D-related developmental and epileptic encephalopathy, a condition associated with developmental delay, intractable seizures, abnormal muscle tone, movement disorders and ASD (*43, 63–65*). NMDAR antagonists such as memantine and ketamine have proven therapeutically useful for treatment of epileptic encephalopathies associated with increased NMDAR function (*43*). In addition to hypotonia and ASD, two individuals carrying *KDM5B* variants show epileptic spasms and generalized seizures (*24, 66*). Given this functional association with NMDARs, future studies assessing the effects of *Kdm5b* mutation on excitatory neurotransmission will be of interest, especially if these can be combined with NMDAR antagonist treatment. Extensive electrophysiological and cell type time-course analyses will be required to characterise the exact phenotypes being rescued by memantine treatment, but our study sets the scene for such further mechanistic work, as well as further exploration of NMDAR antagonists as a therapeutic possibility for individuals with *KDM5B* loss-of-function variants.

Memantine treatment has been evaluated in individuals with ASD, with varied effects, not surprising given that both increased and reduced NMDAR function has been implicated in ASD (*28, 29, 67–71*). The present study, together with studies on *Shank2*-deficient and VPA rodent models (*32, 35*), suggest that treatment with NMDAR antagonists is likely to benefit ASD subtypes associated with increase NMDAR function.

In addition to *Grin2d*, the other differentially-expressed ASD-associated gene worth noting was *Mcm6*. *Mcm6* was the most downregulated gene in *Kdm5b* mutants. De novo *MCM6* variants were first detected in ASD probands by Iossifov et al. (*12*), with further reports by Takata et al. (*40*). Subsequently, missense mutations in *MCM6* were associated with neurodevelopmental phenotypes that include ASD, delayed speech development and epilepsy by Smits et al. (*72*). MCM6 is important for primary ciliogenesis and cell proliferation (*72, 73*), two crucial processes during brain development. Interestingly, previous studies indicate that, in the adult mouse brain, KDM5B is expressed in the neurogenic niches at the subventricular zone (SVZ) and dentate gyrus. The reduction of *Kdm5b* expression led to altered adult neurosphere formation and growth, together with increased migration and differentiation (*74*).

Limitations of our study include the difficulty in directly assessing the contribution of H3K4me3 demethylase deficiency and reduced protein levels on the observed phenotypes. The *Kdm5b^ΔARID^* allele represents a hypomorphic allele that lacks demethylase activity as well as an apparent reduction in steady-state protein levels. It is difficult to distinguish between effects from the reduction in KDM5B protein levels and the lack of demethylase activity. Regardless, we do find a robust upregulation of steady-state H3K4me3 levels in the neocortex, indicating that H3K4me3 demethylase activity is indeed affected in these mice.

In conclusion, we have found that a deficiency in the KDM5B demethylase results in ASD-associated behavioural phenotypes in mice. *KDM5B* deficiency was associated with increased levels of H3K4me3, demonstrating that KDM5B functions as a H3K4me3 demethylase in the developing brain. These alterations affected gene expression, with the majority of differentially expressed genes upregulated, confirming that KDM5B acts primarily as a transcriptional repressor. Finally, we provide evidence to suggest that abnormal elevation of NMDAR2D-containing NMDAR signalling in the postnatal brain, presumably via upregulation of the *Grin2d* gene, is responsible for the socio-communicative deficits in these mice. These observations functionally link *KDM5B* deficiency with childhood neurodevelopmental disorders caused by NMDAR gain of function mutations and suggest a potential therapy to alleviate some of the behavioural dysfunction associated with *KDM5B* deficiency.

## Materials and Methods

### Animals

The *Kdm5b^ΔARID^* mouse line has been described (*26*). In this mouse line *Kdm5b* exons 2, 3 and 4 were removed, leading to the deletion of the entire ARID domain and the truncation of the carboxyl end of the JmJ domain. Mice were maintained in a C57BL/6J background, and experimental animals were produced by *Kdm5b*^+/ΔARID^ x *Kdm5b*^+/ΔARID^ intercrosses. Animals were housed in ventilated cages (37x20x16cm, Techniplas UK Ltd, Leicester, UK) with ad libitum access to water and food (Labdiet PIcolab rodent irradiated, #5R53) and kept at 19–22°C and 40–60% humidity, under a 12:12-h light/dark cycle. The cages contained sawdust (Lignocel wood fiber) and nesting material. A maximum of five animals were housed in the same cage. Our study examined male and female animals, and similar findings are reported for both sexes. All animal procedures were approved by King’s College London AWERB and the UK Home Office (Home Office Project licences P8DC5B496 and PP6246123).

### Genotyping of mice

Genotyping was performed by extracting genomic DNA from ear notches using Proteinase K digestion or the HotSHOT method (*75*) The following primer pairs to detect the WT and the mutant alleles were used: WT forward 5’- CCTTAGACGCAGACAGCACA-3’, WT reverse 5’-CGTGTTTGGGCCTAAATGTC-3’, KDM5B-ΔARID forward 5’-TGCTCCTGCCGAGAAAGTATCC-3’ and KDM5B-ΔARID reverse 5’- CCACCCCCCAGAATAGAATGA-3’. Thermal cycles for the genotyping reactions were as follows: 95°C, 2 min; 35x (95°C, 15 sec; 64°C, 15 sec; 72°C, 15 sec); 72, 12min.

### Western blots

#### KDM5B and Histone 3 western blots

Brain cortices and hippocampi were dissected from postnatal day 5 pups and whole cell protein prepared by lysing in 8M urea, 1% CHAPS, 50mM Tris (pH 7.9) lysis buffer containing protease inhibitors. Samples were rotated for 30min at 4°C and then centrifuged for 60min to remove DNA. Supernatant was stored at -80°C. All reagents and machinery were obtained from BioRad unless stated otherwise. Samples were prepared with Laemmli buffer containing 10% β-mercaptoethanol and resolved with Mini-PROTEAN pre-cast polyacrylamide gels (7.5% and 4-15% gels to analyse proteins with high and medium/low molecular weights, respectively) and Tris/Glycine/SDS buffer. Proteins were transferred to a nitrocellulose membrane with the TransBlot turbo system. Membranes were blocked with 5% non-fat dry milk (Cell Signaling Technologies) or 5% BSA (Sigma) diluted in tris-buffered saline containing 0.1% Tween-20 (TBS-T). Primary antibodies were diluted in TBS-T containing 5% BSA. Primary antibodies were incubated overnight at 4°C in 5% BSA in TBS-T. Membranes were incubated with secondary antibodies diluted in TBS-T containing 5% non-fat dry milk for 1h at room temperature. Proteins were detected with Clarity ECL reagent and membranes were imaged using the Chemidoc system. Densitometric analyses were performed with ImageJ software (NIH). The following antibodies were used (1:1000 dilution unless stated otherwise): anti-KDM5B (Abcam, #181089), anti-H3K4me3 (Cell Signaling Technologies, #9751), anti-total H3 (Cell Signaling Technologies, #9715), HSP90 (Santa Cruz Biotechnologies, sc13119), actin (1:5000, Abcam, #ab8227), goat anti-rabbit and anti-mouse HRP secondary antibodies (1:5000; Thermo Fisher Scientific, #31460 and Proteintech, #SA00001-1, respectively).

#### NMDAR2D western blots

Brain cortices were dissected from postnatal day 5 pups and synaptosomes were isolated using Syn-PER™ Synaptic Protein Extraction Reagent (ThermoFisher) following the manufacturer’s protocol. The synaptosome pellet was resuspended in RIPA buffer containing protease inhibitors and the protein concentration was quantified using a BCA assay (ThermoFisher). Laemmli buffer (Bio-rad) with DDT was added to 15µg of each sample and loaded into a 4–20% precast polyacrylamide gel (Bio-Rad, 4561095). The gel was electrophoresed for 10 minutes at 50V then 1 hour at 120V in Tris-Glycine SDS buffer (Bio-rad). The gel and nitrocellulose membrane were then equilibrised in transfer buffer (Bio-Rad) for 10 minutes before being transferred for 1 hour at 80V in the transfer buffer. The membranes were then stained using Ponceau S (Merck) to check for protein transfer and then blocked in PBS-T containing 5% non-fat milk buffer. The membrane was then incubated with the primary antibodies overnight at 4 °C, and then repeatedly washed with PBS-T before incubation in secondary antibodies for GRIN2D (Merck, MAB5578, 1:500) and B-actin (Merck, A5316, 1:5000) overnight at 4 °C. The membranes were then repeatedly washed with PBS-T before incubating them in Dylight™ 800 secondary antibody (ThermoFisher, SA5-10164, 1:5000) for 1 hour in the dark at room temperature. The membranes were then washed three times and imaged and analysed on the LICOR Odyessey CLx machine.

Uncropped full scans of all the western blot membranes can be found in Supplementary Figure 4.

### Behaviour

Different batches of mice were used for recording pup ultrasonic vocalisations (USVs), for adult behaviour testing and for the treatment with memantine (see Fig 2A and Fig 5C). Adult tests were carried out in the following order: open field, elevated plus maze, reciprocal social tests, marble burying, 3 chamber social approach, and rotarod as previously described (*53, 76*).

Behavioural experiments were conducted between 8:00 and 18:30 under standard room lighting conditions unless stated otherwise. Cages were changed every other week but never less than 24h before the day of testing. Behaviours were tracked using Ethovision (Noldus Information Technologies bv, Wageningen, The Netherlands). After each trial of a specific test, boli and urine were removed, and the test area was cleaned with 1% Anistel® solution (high level surface disinfectant, Trisel Solution Ltd, Cambridgeshire, UK) to remove any odours. The experiments were blinded and randomized by blocks of mice. Littermates were used as controls with multiple litters examined per experiment.

#### 3 chamber social approach

Mice were tested individually in the three-chamber box (20x40.5x22 cm) over three trials. Dividing walls are made from clear perplex, with small openings (10cm width, 5cm height) that allow access into each chamber. Light intensity was 10 lux. During the first trial, mice were habituated to the empty box. In the next trial, an unfamiliar sex-matched juvenile conspecific mouse and an unfamiliar mouse-like object were placed in a wire containment cup on either the left or right chamber (counterbalanced), leaving the middle chamber empty. All trials lasted 10 minutes. The movements of the test mice were tracked during the trials with Ethovision. Conspecific mice were housed in a separate room to the test mice to ensure unfamiliarity.

#### Social interaction

A standard housing cage was used as the test area. Conspecific mice were housed in a separate room to the test mice to ensure unfamiliarity. Test mice were habituated to the test area for one hour with a 1cm layer of clean sawdust, dimly lit (10 lux), with an acrylic lid. Mice did not have access to food and water during the testing sessions. After the habituation period, a same-sex conspecific mouse was placed into the test arena for 5 minutes. If aggressive behaviours were displayed by either mice for more than 10 seconds, the test was stopped. The following investigative behaviours were scored: i) social sniffing: sniffing of the body above the shoulders, ii) anogenital sniffing: sniffing of the anogenital region including the tail and iii) following behaviour: where the test mouse moves in close proximity to the conspecific without making direct contact with them. No aggressive behaviours were observed during testing.

#### Ultrasonic vocalisations (USVs)

Ultrasonic vocalizations (USVs) were recorded in pups across 3-minute sessions in response to social separation from the mother and siblings at postnatal days (PND) 2, 4, 6, 8 and 12 as described (*76*). The tattooing for early identification of the mice was carried out on PND 2. Tattooing (Animal tattoo ink green paste, Ketchum #KI1471) for identification was carried out by inserting the ink subcutaneously through a 0.3 mm hypodermic needle into the center of the paw. Tattoing was carried out at PND2 in the experiment described in Figure 2, and at PND1 for the rescue experiment. USV testing as in Fig.2 was performed on a batch of 22 mice (Males: 3 WT and 6 mutant; Females: 7 WT and 6 mutant). The experiment included in Fig. 5 was performed on a batch of 15 mice (4 male and 3 female saline and 5 male 2 female memantine). During testing, each pup was transferred from its homecage to an empty glass container in a sound-attenuating box placed in a different room. Lights were turned off during the USVs recording. An Ultrasound Microphone (Avisoft UltraSoundGate condenser microphone capsule CM16, Avisoft Bioacoustics, Berlin, Germany), sensitive to frequencies of 10 to 180 kHz, was placed through a hole in the middle of the cover of the sound-attenuating box, about 20 cm above the pup in its glass’s container. USVs were recorded using the Avisoft Recorder software (Version 3.2). The glass container was cleaned with 10% ethanol between pups and with 70% ethanol between litters. For acoustical analysis, recordings were transferred to Avisoft SASLab Pro (Version 4.40) and a fast Fourier transformation (FFT) was conducted. Spectrograms were generated at a frequency resolution of 488 Hz and a time resolution of 1 ms.

#### Marble burying

Repetitive digging behaviour was measured in a test room dimly lit (10-20 lux). Twelve blue glass marbles were placed in a symmetrical 4x3 grid on top of 5 cm deep sawdust bedding in a clean empty cage (37x20x16 cm). Test time was 50 minutes and the number of marbles buried at least ¾ of their height was recorded and counted live at 1 minute intervals.

#### Open field

The circular open field arena was made of clear acrylic with a grey base, with internal dimensions of 40cm diameter and 40cm high. Light intensity inside the arena was 10 lux. Two virtual areas were drawn on Ethovision: a centre zone, 20cm diameter, and a ring 5 cm thick around the perimeter of the arena was defined as the outer zone. Test mice were placed in the open field facing an outer wall to begin the test. Its locomotor activity was tracked for 5 minutes. Total distance moved in the outer zone was used as a measure of locomotive activity level.

#### Elevated plus maze (EPM)

The EPM was made of black Perspex and consisted of four arms (30 x 5 cm). The two opposing closed arms were enclosed by 15 cm high walls on each side and ends. The two opposing open arms were open, as well as the centre platform (5 x 5 cm). The maze was elevated 40 cm above ground. Light intensity was 100 lux on the open arms and 10 lux on the closed arms. The number of entries onto, time spent on, and latency to enter the closed and open arms were manually scored. An arm entry is defined when all 4 paws were located inside the arm. Mice were placed in the centre platform of the EPM facing a closed arm to start the 5 minutes test.

#### Rotarod

Mice were tested in 7 trials over one day on the rotarod. The rod accelerated between 10 to 50 rpm over 300 seconds, and the latency to fall was recorded. Mice were allowed to rest for 5 minutes in their home cage between each trial. If an animal griped to the rod for two consecutive revolutions without walking, it was recorded as a fall.

### Memantine treatment

A memantine (Cayman Chemical Company, #14184) solution was prepared in saline. A fresh solution was prepared for every administration day. 10µl/g of saline or memantine solution were administered intraperitoneally 30 minutes before recording USVs or running the marble burying and accelerating rotarod tests. Doses were 5.6 mg/kg for USVs (*77*) and 10 mg/kg for the marble burying and accelerating rotarod tests (*78, 79*).

### MRI

#### Perfusions

Mice were anesthetized with pentobarbital and intracardially perfused with 30mL of 0.1M PBS containing 0.05 U/mL heparin (Sigma) and 2mM ProHance (Bracco Diagnostics, a Gadolinium contrast agent) followed by 30 mL of 4% paraformaldehyde (PFA) containing 2mM ProHance (*53, 80*). Perfusions were performed with a minipump at a rate of approximately 1 mL/min. After perfusion, mice were decapitated and the skin, lower jaw, ears, and the cartilaginous nose tip were removed. The brain and remaining skull structures were incubated in 4% PFA + 2mM ProHance overnight at 4°C then transferred to 0.1M PBS containing 2mM ProHance and 0.02% sodium azide for at least 1 month prior to MRI scanning (*81*)

#### Magnetic Resonance Imaging – Ex Vivo

A 7-Tesla 306 mm horizontal bore magnet (BioSpec 70/30 USR, Bruker, Ettlingen, Germany) with a ParaVision 6.0.1 console was used to image brains in skulls. Eight samples were imaged in parallel using a custom-built 8-coil solenoid array. To acquire anatomical images, the following scan parameters were used: T2-weighted 3D fast spin echo (FSE) sequence with a cylindrical acquisition of k-space, TR/TE/ETL = 350 ms/12 ms/6, TEeff = 30ms, four effective averages, FOV/matrix-size = 20.2 × 20.2 × 25.2 mm / 504 × 504 × 630, total-imaging-time = 13.2 h. The resulting anatomical images had an isotropic resolution of 40µm voxels.

#### Imaging Registration and Analysis

Image registration is necessary for quantifying the anatomical differences across images. The registration consisted of both linear (rigid then affine) transformations and non-linear transformations. These registrations were performed with a combination of mni_autoreg tools (Collins 1994) and ANTS (advanced normalization tools) (*82*) After registration, all scans were resampled with the appropriate transform and averaged to create a population atlas representing the average anatomy of the study sample. The result of these registrations were deformation fields that transform images to a consensus average. Therefore, these deformations fields quantify anatomical differences between images. As detailed in previous studies (*83, 84*), the Jacobian determinants of the deformation fields were computed and analyzed to measure the volume differences between subjects at every voxel. A pre-existing classified MRI atlas was warped onto the population atlas (containing 282 different segmented structures encompassing cortical lobes, large white matter structures such as the corpus callosum, ventricles, cerebellum, brain stem, and olfactory bulbs to compute the volume of brain structures in all the input images (*85–89*). A linear model with a genotype predictor was used to assess significance. The model was either fit to the volume of every structure independently (structure-wise statistics) of fit to every voxel independently (voxel-wise statistics), and multiple comparisons in this study were controlled for using the False Discovery Rate (*90*).

### RNA sequencing

mRNA quality was analysed using Agilent Total RNA 6000 Pico on a Bioanalyser (Agilent, 2100). Pair-end sequencing (150bp read length) was performed on the Illumina HiSeq 6000 platform. Further data analyses were performed using the Galaxy Europe server (https://usegalaxy.eu). Quality of the raw data was checked using FastQC version 0.11.9. Reads were aligned to the mouse genome (mm10) using RNA STAR Galaxy Version 2.7.8a, and aligned reads were counted using FeatureCounts Galaxy version 2.0.1. Differential expression analyses were performed using DESeq2 Galaxy version 2.11.40.7. Heatmaps were generated with the R package pheatmap. Volcano plots were generated in GraphPad Prism 9.4.1. Gene ontology analyses were conducted using g:Profiler (https://biit.cs.ut.ee/gprofiler/gost) (*91*) on the DEGs (FDR < 0.05), where GO biological processes of size 0-1500 were checked. The applied threshold was “Benjamini Hochberg FDR < 0.05”. Venn diagrams were created to show the extent of overlap between heterozygous and homozygous DEGs (FDR<0.05), and ASD-associated genes obtained from the SFARI human Gene database (https://gene.sfari.org/database/human-gene/; accessed September 2022) or ID-associated genes obtained from the Genomics England PanelApp Intelectual disability database (https://panelapp.genomicsengland.co.uk/panels/285/, version 4.42, accessed December 2022).

### qRT-PCR analysis

cDNA was synthesised from 200-400ng RNA with UltraScript 2.0 cDNA Synthesis kit (PCR Biosystems) according to the manufacturer’s instructions. qRT-PCRs were performed on a BioRad CFX384 using qPCRBIO SyGreen Mix Lo-ROX (PCR Biosystems). See Supplementary table 11 for primer sequences. Relative expression levels were calculated using the 2^-ΔΔCT^ method and *Hprt* was used as endogenous control gene (Sup. Fig. 3).

### Statistics

Data are reported as mean ± SEM. Graphs show individual data points. Normal distribution was tested with d’Agostino and Pearson omnibus, Shapiro-Wilk and Kolmogorov-Smirnov normality tests. If the test was passed, statistical analysis was performed using parametric statistical analyses. Before pairs of comparisons, we performed the F test to compare variances. Statistical analyses were performed using the unpaired two-sided Student-s t test and the three- and two-way ANOVA with the appropriate post hoc tests as indicated in the figure legends. T test with Welch’s correction was applied when variances were unequal. Significant p values (p<0.05) are reported in the results section and/or figure legends provide details of relevant statistical parameters, including group sizes. Statistical analyses were performed with GraphPad Prism (version 9.4.1). Experiments in this study were blinded and in vivo studies were randomized by blocks of animals.

## Acknowledgements

We thank animal husbandry staff at KCL for expert animal care. We thank our laboratory colleagues for comments on the manuscript. The Galaxy server that was used for some calculations is in part funded by Collaborative Research Centre 992 Medical Epigenetics (DFG grant SFB 992/1 2012) and German Federal Ministry of Education and Research (BMBF grants 031 A538A/A538C RBC, 031L0101B/031L0101C de.NBI-epi, 031L0106 de.STAIR (de.NBI)).

## Author contributions

MAB and LPS conceived the study and designed the experiments. LPS, SB, AC, TG, SS, EH and JE performed the experiments and analysed the data. LPS and MA performed bioinformatic analyses. JTP provided the *Kdm5b*-deficient mouse line and associated expertise. DC provided advice and assistance with memantine experiments. NC, LCA, JL, MLS, KPG, CF and MAB oversaw the research and analysis. LPS and MAB wrote the manuscript with input from co-authors.

## Data availability

All data needed to evaluate the conclusions in the paper are available in the main text and/or the supplementary materials. RNAseq raw data has been deposited at the Gene Expression Omnibus (GEO) archive (GSE262555) and will be made freely available upon publication.

---

**Fig. S1.**
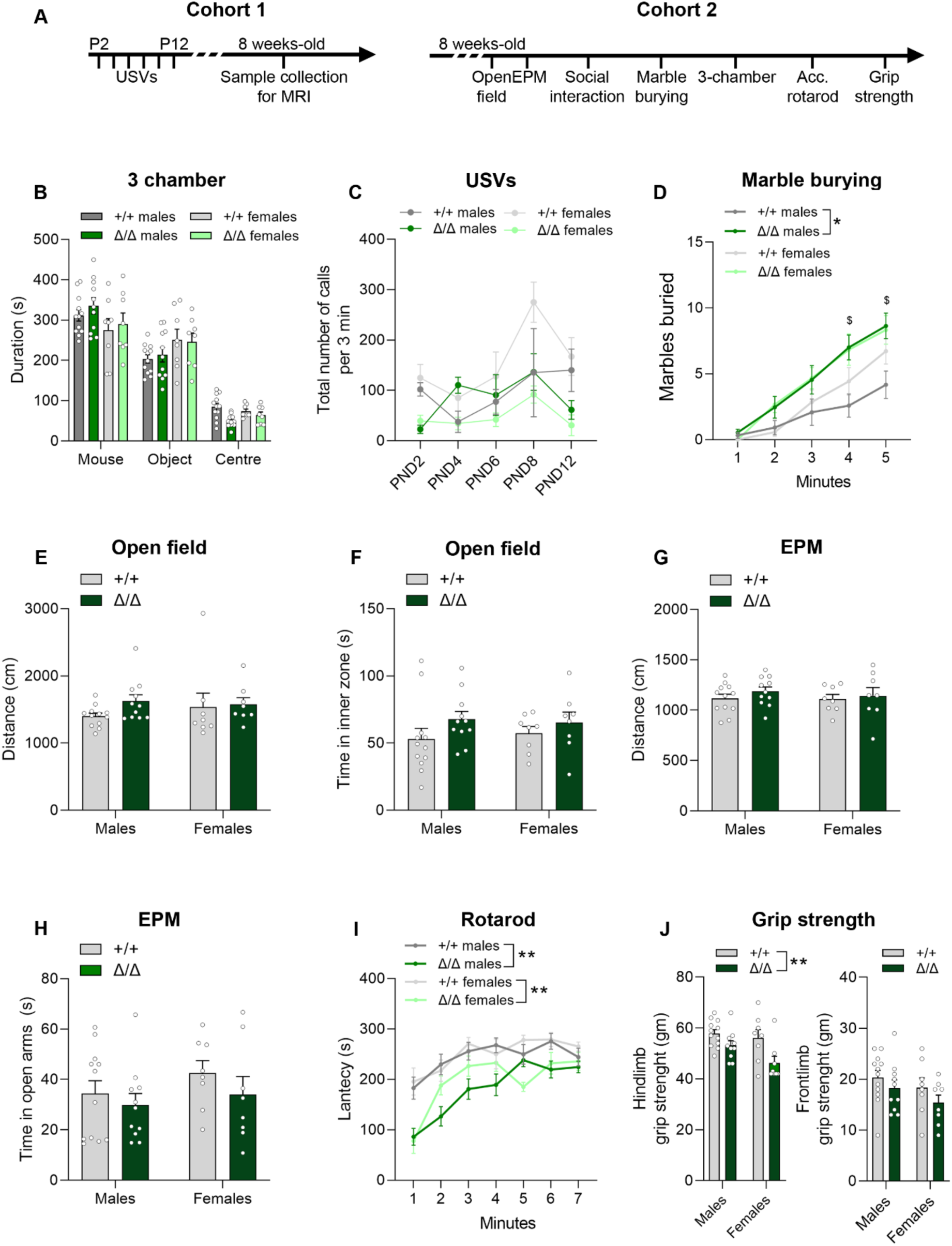
Behavioural assessment of *Kdm5b* ΔARID homozygous mice. Behavioural assessment of a cohort of adult mice (A, C-I) (+/+ n=11-12 males and 7-8 females; Δ/Δ n=10-11 males and 6-8 females) and pups (B) (+/+ n=3 males and 7 females; Δ/Δ n=6 males and 6 females). A) Experimental time-course. Note that different cohorts were used for experiments with neonatal and adult mice. B) Time spent, in seconds, in each chamber in the 3 chamber sociability test, for male and female animals. Three-way ANOVA sex effect: F_(1,105)_=2.49e-005, p=0.9960. C) Total number of ultrasonic vocalisations (USVs) during 3 minutes testing sessions on the indicated postnatal days. Three-way ANOVA sex effect: F_(1,18)_=0.3555, p=0.5584. D) Number of marbles buried during a 5 minutes test. Three-way ANOVA sex effect: F_(1,3160)_=0.9021, p=0.3437. $ p<0.05 mutant male versus WT male. E) Distance travelled in the outer area of an open field arena during a 5 minute test. Two-way ANOVA sex effect: F_(1,35)_=0.1520, p=0.6990. F) Time spent in the inner zone of the arena during the open field test. Two-way ANOVA sex effect: F_(1,35)_=0.01616, p=0.8996. G,H) Distance travelled (F) and time spent (G) in in the open arms of the elevated plus maze (EPM) is shown. Two-way ANOVA sex effect: F_(1,35)_=0.2334, p=0.6320 and F_(1,35)_=1.263, p=0.2688, respectively. I) Mean latency of mice to fall from the rotarod during 7 trials in one day. Three-way ANOVA sex effect: F_(1,35)_=0.7865, p=0.3812. J) Hind and frontlimb grip strength. Data is shown as mean ±SEM and was analysed with Two-way ANOVA (A, D-G, I) followed by Tukey’s multiple comparisons test or repeated measures ANOVA (B, C, H) followed by Sidak’s post hoc test. Sex differences were assessed with three-way ANOVA (A-C, H). \**P*<0.05, \*\**P*<0.01.

**Fig. S2.**
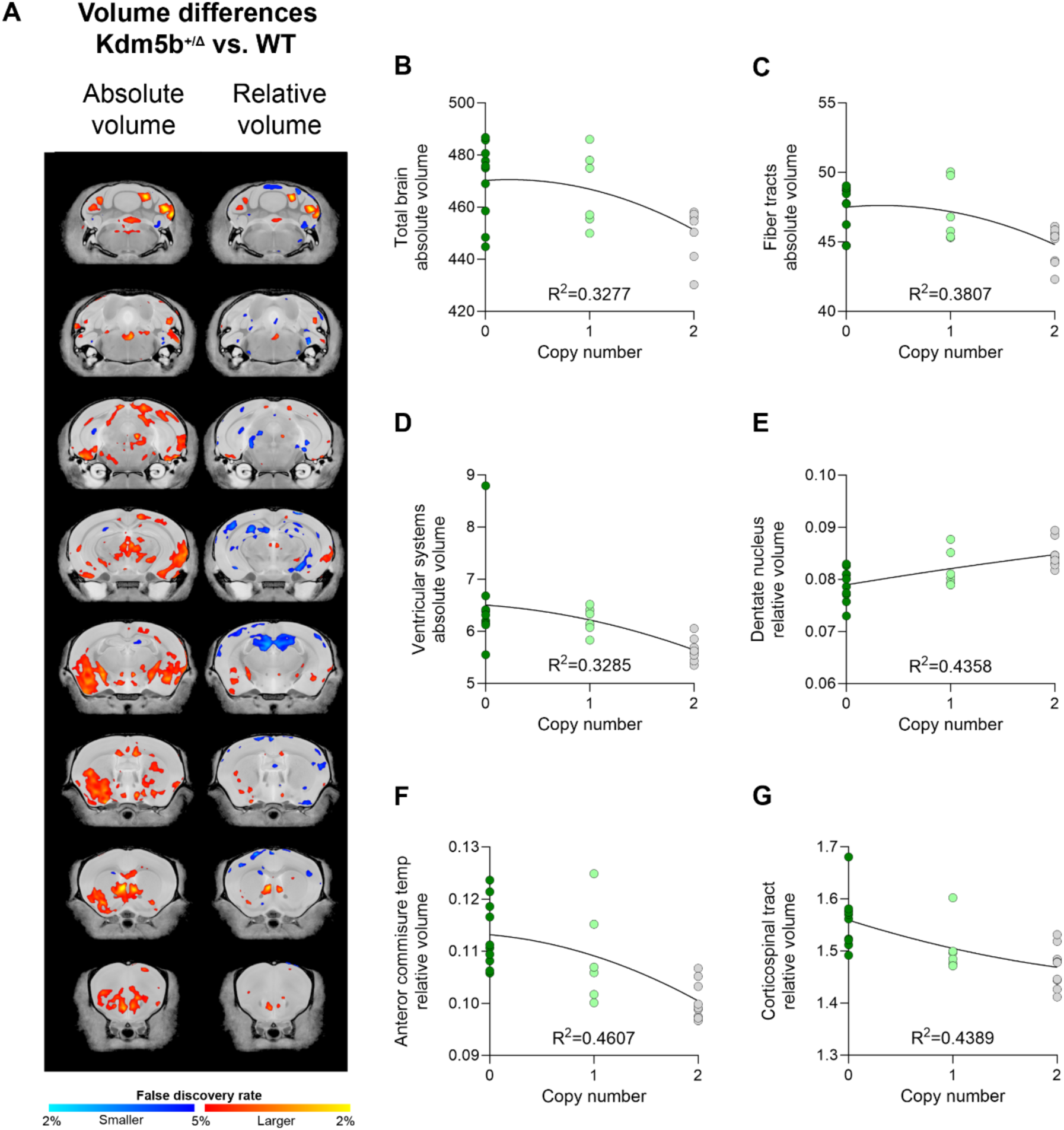
A) Voxel-wise differences in volume (absolute, left; relative, right) between wild-type and *Kdm5b*^+/Δ^ mutant littermates. B-G) Nonlinear (polynomial) correlation analysis between absolute and relative volumes and gene copy number. Each point represents an individual animal.

**Fig. S3.**
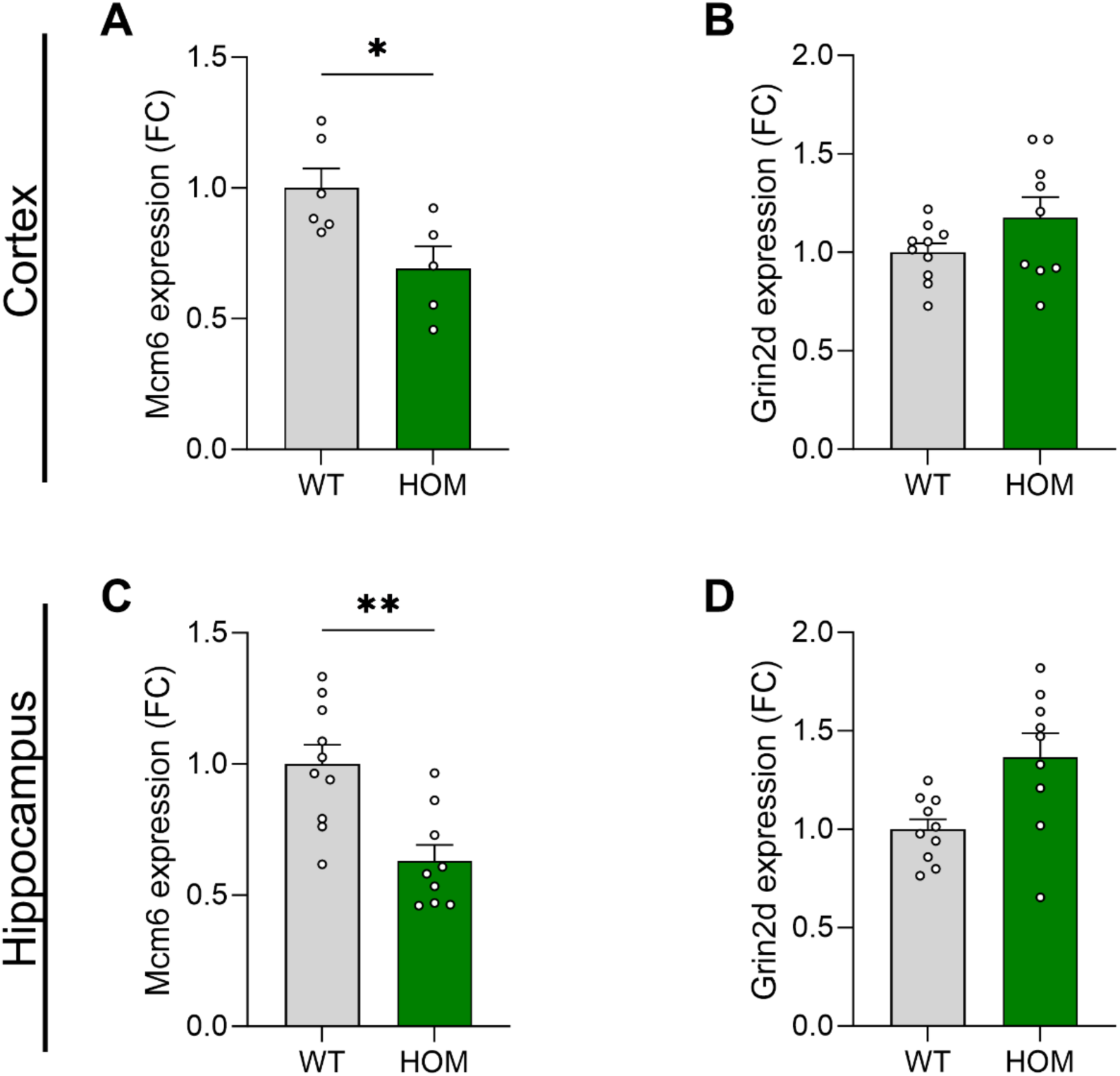
qRT-PCR analysis of *Mcm6* and *Grin2d*, relative to *Hprt*, from total RNA extracted from the cortex (A, B) or hippocampus (C, D) from P5 mouse brains. Student’s t-test. \**P*<0.05, \*\**P*<0.01. N=8 males +/+, 8 males Δ/Δ, 6 females +/+ and 5 females Δ/Δ.

**Fig. S4.**
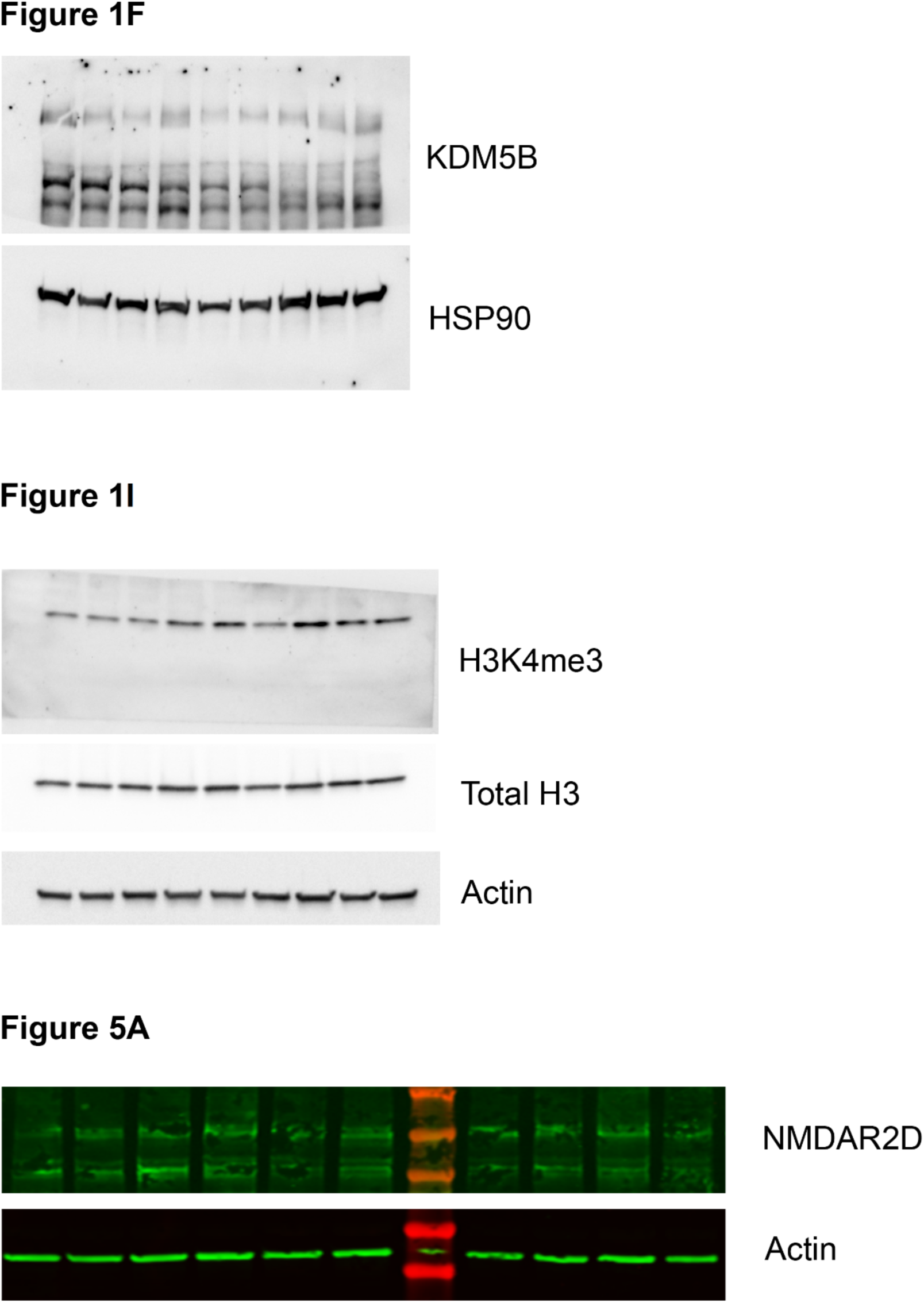
Uncropped western blot full scans for the corresponding blots on Figures 1 and 5.

